# Towards a comparative framework of demographic resilience

**DOI:** 10.1101/2020.01.31.928721

**Authors:** Pol Capdevila, Iain Stott, Maria Beger, Roberto Salguero-Gómez

**Affiliations:** Department of Zoology, University of Oxford, 11a Mansfield Rd, Oxford, OX1, 3SZ, UK; Department of Biology, Interdisciplinary Centre on Population Dynamics (CPop), University of Southern Denmark, 5230 Odense M, Denmark; School of Life Sciences, University of Lincoln, Brayford Pool, Lincoln, LN6 7TS, UK; School of Biology, Faculty of Biological Sciences, University of Leeds, Leeds, UK; Centre for Biodiversity and Conservation Science, University of Queensland, St Lucia 4071 QLD, Australia; Evolutionary Demography laboratory, Max Plank Institute for Demographic Research, Konrad Zuße Straße 1, Rostock 18057, Germany

**Keywords:** Global Change, Life History Strategies, Regime Shifts, Stability, Stage-Structured Population Model

## Abstract

In times of global biodiversity crisis, developing tools to define, quantify, compare and predict ecological resilience is essential for understanding species’ responses to global change. Disparate interpretations of ecological resilience have, however, hampered the development of a common currency to quantify and compare resilience across natural systems. Most frameworks of study have focused on upper levels of biological organisation, especially ecosystems or communities, which adds layers of complication to measuring resilience with empirical data. To overcome such limitations, we suggest quantifying resilience using demographic data. Surprisingly, a quantifiable definition of resilience does not exist at the demographic level. Here, we present a framework of demographic resilience with a set of metrics that are comparable across species, and facilitate cost-effective management decisions.

## Resilience as a key concept in ecology and conservation

Contemporary global change is increasingly eroding the natural resources we depend on [1,2], and understanding how ecological systems withstand these disturbances is a major challenge [3–5]. “Resilience” is a key concept describing natural systems’ abilities to handle disturbances [6]. Indeed, international environmental policy objectives, including the UN Sustainable Development Goals [7] and Aichi Targets [8], specifically include preserving resilience as a key objective.

Resilience describes the ability of a system to recover from and persist after a disturbance [6]. However, translating this concept to quantifiable metrics is challenging due to the complex nature of ecological systems [9], generating multiple debates over the past decades regarding the definition, meaning and application of resilience [10–12] (Box 1). Discrepancies between approaches mean both theoretical and empirical works lack parity between the primary components of resilience studied, rendering comparisons challenging if not impossible. These limitations ultimately prevent ecologists from applying resilience-based solutions to real-world problems (e.g. see [13]). Developing a unifying framework with comparable definitions and quantifications across different ecological systems is therefore an urgent task [14], with recent studies advocating tangible and meaningful resilience measures [11,12,14]. Despite populations often being the target of conservation interventions [15], no formal framework exists for defining and quantifying their resilience.

We introduce a framework to define, quantify, and compare resilience across populations and species. The framework utilises classical [21] and recent demographic approaches [17,18] alongside resilience theory [12,14,17,18]. All populations are ruled by demographic processes including rates of survival, development, and reproduction [19] that ultimately determine their temporal dynamics, vulnerability and management [19]. Thus, demographic processes constitute the ideal common currencies to quantify demographic resilience. Such a common currency facilitates comparison of the same resilience metrics across different species or populations.

### Box 1

**The meaning of resilience**

Since its first appearance in the ecological literature in the late 1970s, the study of resilience has attracted a significant amount of attention (Figure I). However, the rate at which research in the area has increased is comparable to the diversity of definitions and different interpretations of resilience. The term resilience was first introduced to ecology by Holling [6], who defined it as “a measure of the persistence of systems and their ability to absorb change and disturbance and still maintain the same relationships between populations or state variables”. Despite Holling’s clear, comprehensive definition, following authors/sub-disciplines interpreted it in different ways [20]. For example, some authors considered resilience as the speed of recovery of a natural system, quantified as the time required to return to equilibrium [21]. In contrast, other authors have measured resilience as the probability of the system to remain above their unstable equilibrium [22]. Consequently, later on, Holling [23] distinguished two types of resilience: engineering and ecological resilience. He defined engineering resilience as the rate or speed of recovery of a system following a shock. Ecological resilience, meanwhile, was described as the magnitude of a disturbance required to trigger a shift between alternative states [6,23]. Such a distinction was made to stress the importance of the existence of multiple stable states in ecological systems [23]. While ecological resilience does account for the existence of multiple stable states, engineering resilience assumes only a single equilibrium point.

Recent evidence, however, shows that resilience can be achieved in different ways [12,24–26]. For example, a natural system may show some opposition to an external disturbance, limiting its displacement from its initial state, showing resistance to change. On the other hand, a system can show low resistance to disturbances, but may have a high ability to come back to its initial state, displaying a fast recovery. Several authors have suggested framing resilience as the result of resistance and recovery [12,14,26], because it can capture the different ways through which natural systems respond to disturbances. Here, we align with the definition of resilience that includes resistance and recovery time as two integral parts of the ecological system.

**Figure I.**
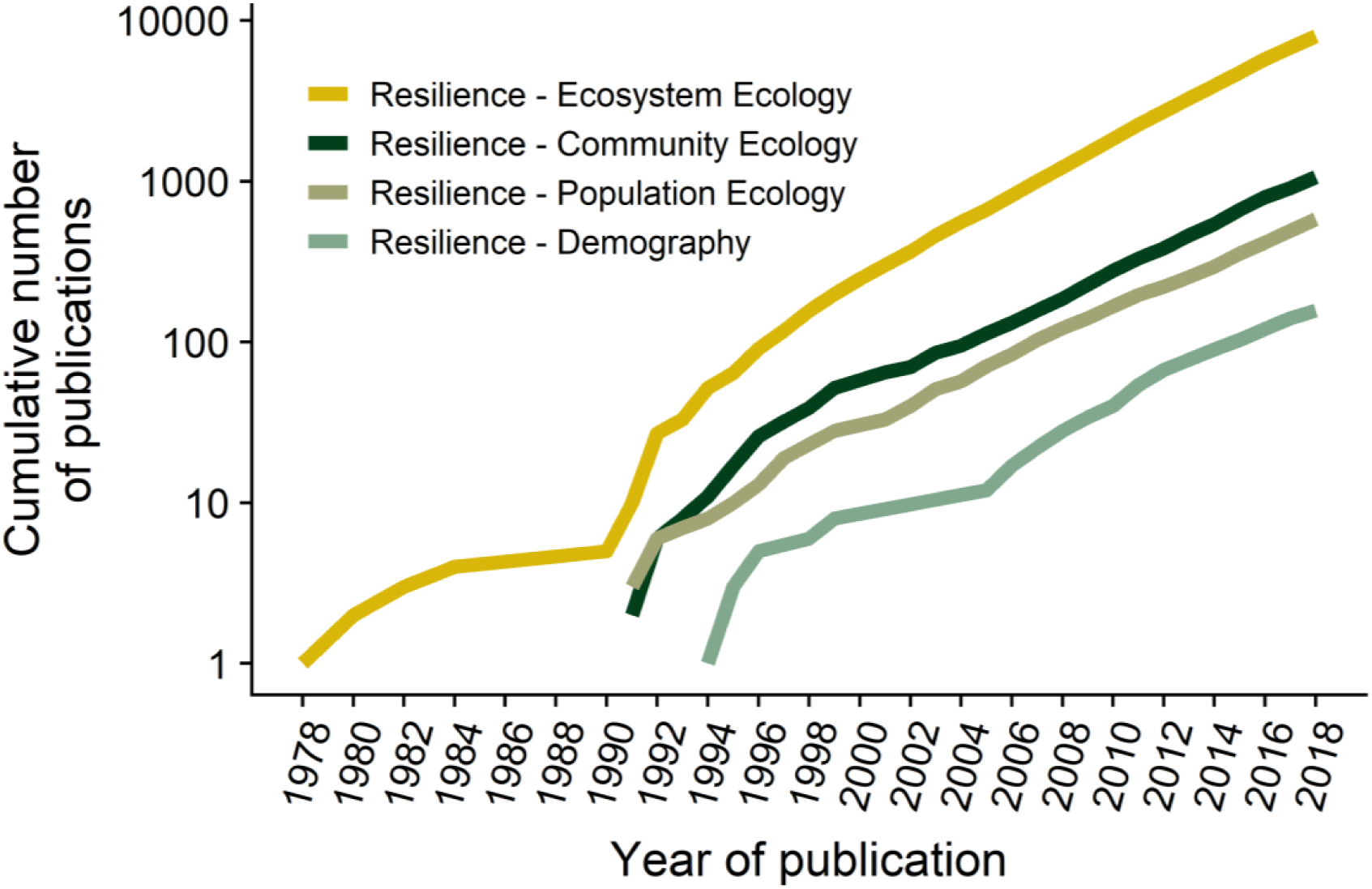
The cumulative number of ecological studies has increased exponentially in the last decades, but less so in lower levels of biological organisation such as physiology or population ecology. The cumulative number of publications (in logarithmic scale) in the Web of Science was obtained with the search criteria: “Resilience * Ecosystem Ecology”, “Resilience * Community Ecology”, “Resilience * Population Ecology” and “Resilience * Demography”, respectively. The literature search was constrained between 1^st^ January 1945 and 31^st^ December 2018.

## Theoretical measurements of resilience

Established resilience theories assume that natural systems can exist in alternative stable states [6], where the forces influencing the system are in balance [6,20,21,22]. When a disturbance displaces the system to an unstable state, these forces usually draw it back to stability. However if a strong disturbance forces the system beyond a domain of attraction, a *tipping point*, the system may transition to an alternative stable state [17,18]. This new system state is characterised by substantially different structures and maintained by processes of *hysteresis* or feed-backs [17,27].

These classical theoretical frameworks have triggered the development of a myriad of resilience indicators [17,18,28]. These indicators are based on the idea of *critical slowing down*, whereby a system approaching a tipping point may exhibit decreasing ability to recover its previous state due to a decline in its resilience [17,28]. Approach to a critical tipping point can be detected with temporal and spatial statistical signatures, such as increased autocorrelation of, or variance in, abundance [18,28]. Such momenta have been identified in different ecosystems [17,18], potentially facilitating anticipation of critical system transitions [29,30].

Detecting approaches to tipping points is debated [13,31], given their limitations related to (*i*) assuming abrupt regime shifts [32], (*ii*) assuming regime shifts exhibit critical slowing down [18,32], and (*iii*) the inability to compare systems with dissimilar properties and/or environments [18,28]. This theoretical framework is further unable to (*iv*) explicitly account for different responses to disturbances for the different species life history strategies [33,34], and (*v*) distinguish population responses prior to collapse [28,35] from responses to disturbance. Such constraints (discussed further in [28,35]) have hampered the use of resilience theory [11,13] in applied ecology and conservation.

## Demographic resilience

A new demographic resilience framework can mutually inform and complement existing community resilience theories. Here, we develop a framework for understanding population resilience, drawing on ideas and terminology from community resilience. A framework for demographic resilience can tackle many challenges associated with community resilience by: relaxing the assumption of systems experiencing regime shifts and tipping points (limitations *i* and *ii*), because it focuses on the responses of the populations to disturbances [16]; allowing comparison of the same fundamental processes (survival, development, and reproduction) across different populations and/or species (*iii*) [19]; accounting for the differences in the life histories (*iv*); and estimating the population responses prior to a collapse (*v*), by quantifying their dynamics [36].

Populations show similar properties to classical community resilience frameworks. Just like communities, populations are structured [37]: as distinct species in a community contribute differently to community dynamics [38], individuals of distinct age, size or developmental stage in a population contribute differently to population dynamics [37]. In a constant environment with unlimited resources, a population will attain a stable structure with a stable long-term growth (or decline) [16,37]. Disturbances typically change population size and structure, displacing it from stable growth (e.g. a fire affects more young rather than old trees [39]). Short-term *transient* growth is faster or slower than stable growth (*amplification* and *attenuation* respectively [16]; Figure 1B).These are respectively generated by a relative over- or under-representation of individuals with high survival and reproduction. Thus, transient dynamics depend on population structure [19,37]. As under-represented individuals are repopulated, the population is drawn back towards stable state over the *transient period*; akin to recovery in classical resilience theory (Figure 1).

**Figure 1.**
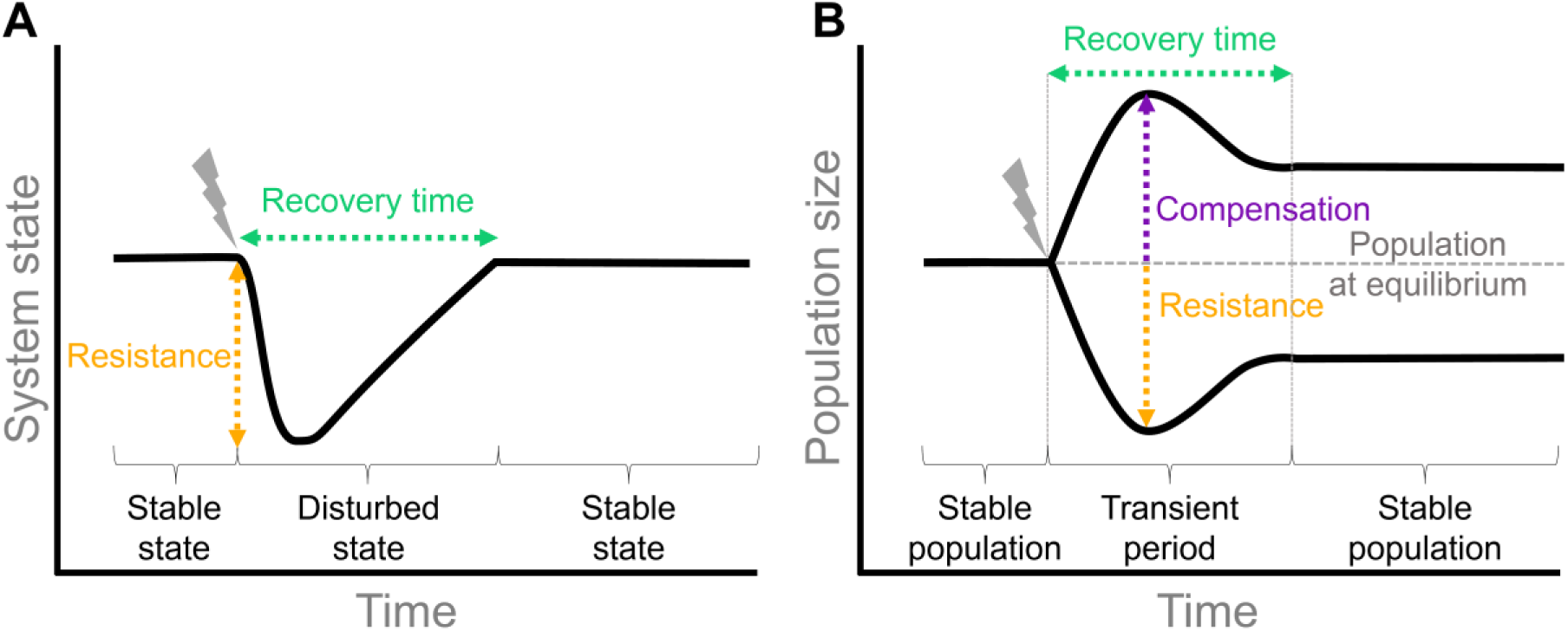
Comparison between disturbance responses and the main components of resilience in communities (A) and populations (B). When translating the population responses to disturbances from classical resilience frameworks, the system state is defined as the population size and the population structure (y axis). After a disturbance, the size of the population change differently according to the stages impacted, creating a range of possible population sizes, and defining the resistance of being disturbed. The time needed to settle to one of the multiple possible stable structures is defined as the recovery time. The decrease of the population after a disturbance is resistance. In demography (B), there is another possible response to disturbance compared to communities (A), which are increases in population size or compensation.

## Measuring and comparing demographic resilience

The extensive quantitative development of population ecology provides a corollary of tools to measure population resilience, overcoming one of the main criticisms of existing resilience frameworks in communities [12,13]. Structured population models facilitate explicit simulations of disturbances impacting different life cycle stages [16,37], and enable calculation of the consequent transient responses. These represent three key components of resilience: demographic ***compensation, resistance***, and ***recovery*** (Figures 1 and 2). We explicitly link each measurement to the dimensions of resilience that it quantifies below (Box 2).

**Figure 2.**
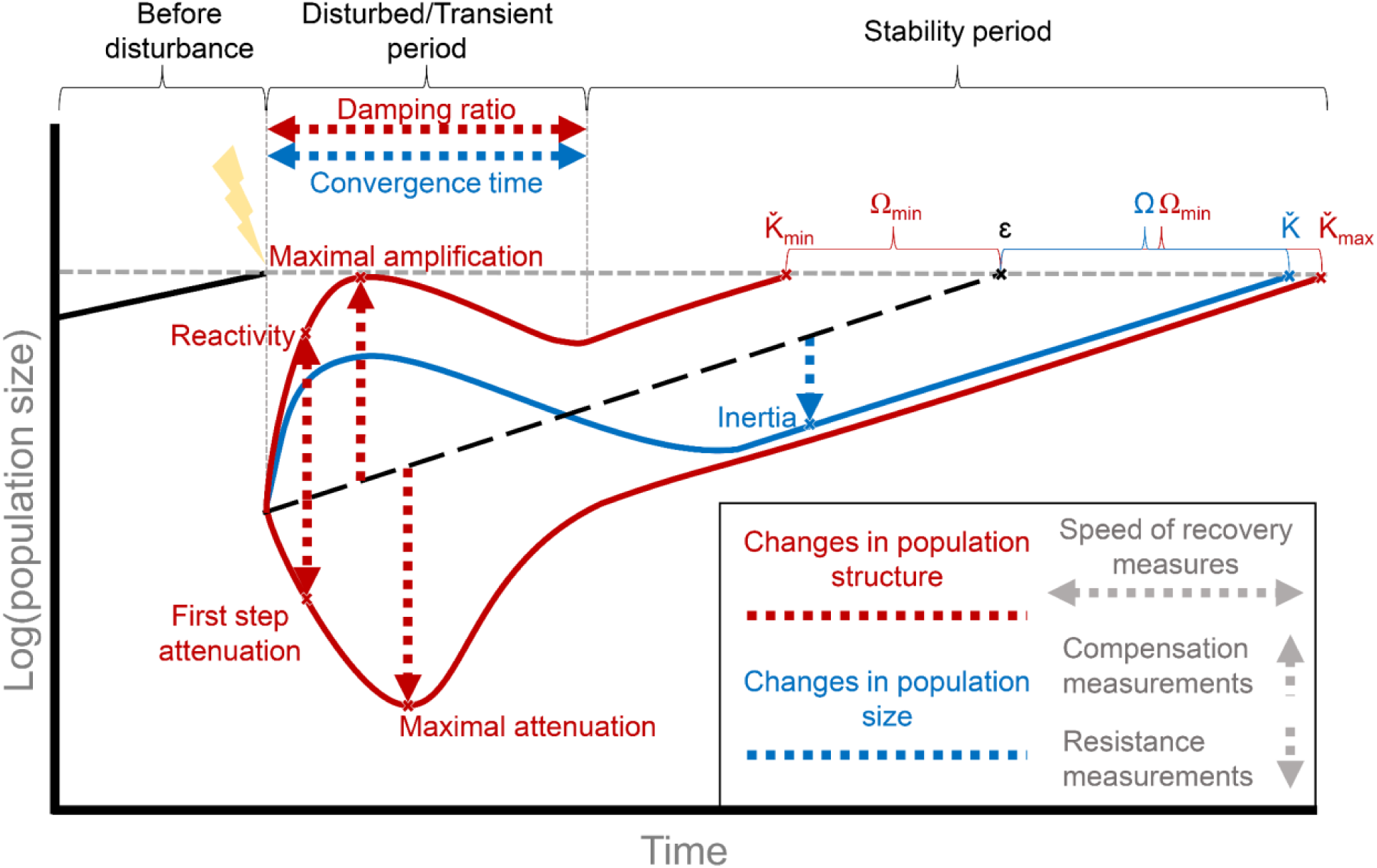
Resilience framework measurements for populations’ responses to disturbances. Example of a population impacted by a disturbance. Before the disturbance, in this example the population is increasing (but could also be decreasing or remain stable). After the disturbance, imbalances in the proportion of individuals at each stage cause population increases or decreases, creating a discrepancy between the actual population size/structure and the one that would exist given stable growth following the disturbance. At the first-time step after the disturbance, the population density increase and decrease are *reactivity* and *first step attenuation*, representing the immediate response of the populations. During the transient period the population depict from stable structure, but the population will tend towards stability. The time elapsed for the population to reach stability can be estimated as the damping ratio or convergence time, measurements of speed of recovery. During this transient period, the highest and the lowest population density after disturbance represent the *maximal amplification* and the *maximal attenuation.* Once reached stability, the disturbance may have created a discrepancy between the initial stable size/structure with the long-term one, with the upper bound measured as *amplification inertia* and the lower bound as *attenuation inertia*. In addition, it is possible to estimate the time required to recover the initial stable population structure has its minimum at 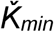 and maximal at 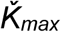. The difference between 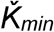 and 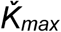 to the structure at the stable population growth *ε*, it is possible to estimate *Ω*_*min*_ and *Ω*_*max*_ to measure of how much time the system will require to reach the initial structure. It is similar for population size, with 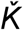 being the time to reach stability and *Ω* being the difference with stable growth.

### Demographic compensation

*Demographic compensation* incorporates *amplifications* in population size after disturbance. Population *amplification*, quantifies population increases following a disturbance (Box 2, Figure 2). We advocate the use of *reactivity, maximal amplification* and *amplification inertia* [16] to estimate changes in population size and structure at various times after a disturbance (Figure 2). Reactivity quantifies the *immediate*, short-term response to a disturbance, maximal amplification is the *highest* density that the population can reach at *any time step*, and inertia measures the *total* displacement of the population on the long-term. Reactivity, therefore, quantifies immediate compensation of a population, maximal amplification measures the overall ability of the population to compensate and inertia quantifies how far away from the stable state the population ends up, as a result of transient dynamics following disturbance.

Demographic compensation is fundamental for understanding population crashes [16], and compensation metrics are of particular interest for management actions targeting potential invasive species [40]. For instance, species showing high population increases after disturbance can be a potential problem for managers, who may wish to adapt their management interventions according to the potential population amplification [15,40]. For that reason, even if not considered in classical views of resilience (e.g. [6]), we advocate including demographic compensation in resilience studies.

### Demographic resistance

If we consider resistance as a measure of the proportion of a variable that changes before/after a disturbance [12,14,26], *demographic resistance* can be estimated by incorporating both population amplification and attenuation. The largest possible amplification and attenuation values, the so-called *transient bounds*, represent the most extreme possible values of transient population size, and together they represent the *transient envelope* (Fig. 1; [16]). A small transient envelope means that population size is robust against disturbance (*i.e.* more resistant), while large transient envelopes indicate that the population is more sensitive to changes in its structure [16,41] (less resistant; see Box 2). As amplification and attenuation are bound asymmetrically ([1, ∞) for amplification; (0, 1) for attenuation [16]), arithmetic rather than geometric differences in growth are more relevant resistance measures. Note that in Box 2 we did not include the transient envelope for maximal amplification and attenuation, given that both can happen at different times. The transient envelope is useful for comparative studies given its intuitive interpretation within and across populations.

Population attenuation bounds can also be used as proxies of population resistance (Figure 2). Similarly to population compensation and for the transient envelope, we suggest using *first step attenuation, minimum attenuation*, and *attenuation inertia* [16] to estimate the potential change in population size and structure after a disturbance (Box 2). First step attenuation quantifies the immediate response to disturbance, maximal attenuation is the lowest density that the population can reach at any time, and attenuation inertia measures the total displacement on the long-term. Consequently, first step attenuation quantifies the magnitude of population decay or lack of resistance, maximal attenuation measures the overall lack of resistance, and inertia quantifies how far away from the stable state the population ends up.

At the community level most works express resistance as a measure of the loss of species after a disturbance or change in community structure [42–44]. Community resistance can be measured as the maximal Euclidean distance between vectors representing a perturbed and an unperturbed community. The higher the Euclidean distance the lower the community resistance, and *vice versa* [9,45], whilst multi-dimensional variables are aspects of the quality and diversity of the community before and after the disturbance [9,45]. We advocate that population resistance can be measured using differences in population size, *i.e*. the sum of the population’s age or stage vector. This approach is in essence the same as that already used for communities, but using a more intuitive means of quantifying the system in state space: the Euclidean distance in communities versus the vector sum for populations.

### Time of recovery

*Time of recovery* is a critical metric of demographic resilience that explicitly considers time. Similar to resistance metrics, there exist a number of metrics to quantify the time required to reach population stability [16]. For populations, the key question is *time of recovery to what?* Stable state, or a desired population size/structure? We propose two measures to describe the time of recovery to population stability after a disturbance: *damping ratio* and *time of convergence* (Box 2). We also propose two metrics to estimate *time to recover population size* and *population structure* (Box 2).

### Speed of recovery to stable state

The damping ratio measures how quickly transient dynamics decay following a disturbance, regardless of the population structure [16]. The larger the damping ratio, the faster the population converges, and the higher the speed of recovery. Importantly, the damping ratio is a dimensionless metric [37], *i.e.* it possesses no units. Thus, damping ratio is useful to *compare* relative time of recovery across populations or species [36]. In contrast, though the time of convergence is similar to the damping ratio, the former is time-stamped, so it can be used both for comparative analyses and to inform managers about the expected post-disturbance recovery times.

### Time of recovery to population size and structure

If the population was not in stability before the disturbance, it is also possible to estimate time required to recover previous the population size and/or the original structure (Figure 2). Because returning times to population size or structure can be measured relative to any desired structure, such metrics can provide useful insights for conservation plans or restoration actions.

At the community level, time of recovery has been sometimes defined as resilience [13,46]. Recovery time has been estimated using a wide variety of measurements, sometimes specific to the study system, such as net primary productivity [47] or biomass [48]. The common denominator is that such metrics are compared between the disturbed and undisturbed communities after certain intervals of time. In the case of empirical studies, such intervals are constrained to the length of the study, and so a full recovery is not always observed [47,48]. In contrast, modelling studies can project the community and measure its recovery at long temporal scales [45].

## Concluding remarks and future perspectives

Our proposed framework extends community resilience [12,14,28,49] to demographic resilience. Demographic resilience allows operationalising and comparing resilience across different species, overcoming two of the main challenges of resilience research. By framing resilience through a population ecologist’s lens, we provide a set of tools that define and enable the quantification of resilience at the population level, and the comparison of resilience across different species.

Demographic resilience opens the door to multiple research venues (see Outstanding Questions). Comparing demographic resistance and recovery across species will allow quantification of differences and commonalities in resilience, and the mechanism by which resilience is achieved. Such information will be crucial for informing conservation science in developing management and conservation actions specific to relevant components of resilience (e.g. estimating the recovery potential of species [15]). Operationalising resilience across species will also enable the exploration of evolutionary questions. For example, by integrating phylogenetic comparative analyses [50] with demographic resilience estimates, one could infer the resilience for data-poor species through closely-related, data-rich species.

Disturbance regimes are important determinants of demographic resilience. Our framework quantifies resilience as a standard property of the life history, across potential disturbances, by quantifying potential outcomes from the changes in the population structure [16]. Specific disturbance regimes can, however, be explored by estimating case-specific transient dynamics where population structure following the disturbance is known [16]. Here, we define disturbance as a sudden event, *i.e.* a pulse of mortality caused by a temporary period of environmental stress altering the natural state of the system (e.g. storm, fire) [51]. However, beyond sudden and fleeting disturbance events, chronic events called *perturbations* that have sustained influence on populations (e.g. global warming, ocean acidification) are also likely to influence population resilience [51]. The adequacy of considering chronic events in a resilience context has been debated [12,52], with some authors considering them to cause a permanent system change, so a return to stability can only be achieved through adaptation [12]. The resistance and recovery framework may not apply in such cases [12], and incorporation of adaptation might be required (e.g. [53,54]).

Because the demography of a species is tightly linked to biological processes taking place at lower and higher levels of organisation, our framework enables exploration of the constituent mechanisms driving resilience. Resilience is an emerging property of complex systems [55], and can be seen as a by-product of the multiple, individual resilience of the subcomponents of the system [56]. Considering that ecological communities are assemblages of populations of interacting species, [42], understanding demographic resilience could provide important insights on how community resilience arises. As individual elements of the community become less resilient, the community will likely be more susceptible to disturbances. The links between demographic resilience and physiological resilience are also likely to provide mechanistic insights on how individual’s resilience scales up into populations and communities. For example, species with fast speeds of recovery are likely to have individuals with strong physiological resilience, because losses of individuals due to disturbances would need a quick repopulation through recruitment and reproduction.

### Box 2

**Transient calculations**

The estimation of transient dynamics can be done in different ways [16]. They can be measured estimating the absolute changes in the population size, which combine the transient rates and the asymptotic. However, the asymptotic effects can be discounted by using a standardised MPM **Â**, by dividing matrix **A** by λ_max._ Also, the population vector **n** can also be standardised 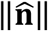 to sum to 1. Such standardisations are highly recommended because they allow fair comparisons among models and then are useful for both conservation and comparative analyses [16].

We present here a compendium of equations to estimate the abovementioned transient metrics.

| Resilience component | Index | Transient calculations | Calculation | Interpretation |
| --- | --- | --- | --- | --- |
| Population compensation | First step population increase | Reactivity | $\ \hat{\mathbf{A}}\ _1$ | The largest population density that can be reached in the first time step after disturbance |
| | Maximal population increase | Maximal amplification | $\bar{\rho}_{\max} = \max_{t>0} (\ \hat{\mathbf{A}}^t\ _1)$ | The largest population density that can be reached at any time after disturbance |
| | Long-term population increase | Amplified inertia | $\frac{v_{max} \ w\ _1}{v^T w}$ | The largest possible long-term population density |
| | Reactivity envelope | Reactivity multiplied by first step attenuation | $\ \hat{A}\ _1 * minCS(\hat{A})$ | The higher the value, the greater the tendency of the population towards amplification over attenuation following disturbance. |
| | Inertia envelope | Amplified multiplied by attenuated inertia | $\frac{v_{max} \ w\ _1}{v^T w} * \frac{v_{min} \ w\ _1}{v^T w}$ | The higher the value, the greater the tendency of the population towards amplification over attenuation in population displacement. |
| Resistance | First step population decrease | First step attenuation | $minCS(\hat{A})$ | The lowest population density that can be reached in the first time step after disturbance |
| | Maximal population increase | Minimum attenuation | $\underline{\rho}_{min} = \min_{t>0} (minCS(\hat{A}^t))$ | The lowest population density that can be reached at any time after disturbance |
| | Long-term population increase | Attenuated inertia | $\frac{v_{min} \ w\ _1}{v^T w}$ | The lowest possible long-term population density |
| | Reactivity envelope | Reactivity divided by first step attenuation | $\ \hat{A}\ _1 / \min CS(\hat{A})$ | The lower the value, the more the population resists changes in size. |
| | Inertia envelope | Amplified divided by attenuated inertia | $\frac{v_{max} \ w\ _1}{v^T w} / \frac{v_{min} \ w\ _1}{v^T w}$ | The higher the value, the greater the displacement of the population from its stability in the long term after disturbance |
| Speed of recovery | Damping ratio | The dominant eigenvalue ($\lambda_1$) divided by the absolute value of the largest subdominant eigenvalue ($\lambda_2$). | $\rho = \lambda_1 / \ \lambda_2\ $ | Dimensionless measure of convergence to stability |
| | Convergence time | The time $t_x$ required for the contribution of $\lambda_1$ to become x times as great as that of $\lambda_2$ | $\log(\rho)/\log(x)$ | Time of convergence to stability |
| | Minimum time to recover initial structure | Time to recover initial population structure at stable growth rate minus the minimum time to recover initial population structure | $\Omega_{min} = \varepsilon - \check{K}_{min}$ | The lower the value the less time required to recover the initial population structure |
| | Maximal time to recover initial structure | Time to recover initial population structure at stable growth rate minus the maximal time to recover initial population size | $\Omega_{max} = \varepsilon - \check{K}_{max}$ | The lower the value the less time required to recover the initial population structure |
| | Time to recover initial population size | Time to recover initial population size at stable growth rate minus the maximal time to recover initial population size | $\Omega = \varepsilon - \check{K}$ | The lower the value the less time required to recover the initial population size |

